# Physiological starvation increases EGF-Ras-MAPK pathway activity during *C. elegans* vulval induction

**DOI:** 10.1101/329573

**Authors:** Stéphanie Grimbert, Amhed Missael Vargas Velazquez, Christian Braendle

## Abstract

Studying how molecular pathways respond to ecologically relevant environmental variation is fundamental to understand organismal development and its evolution. Here we characterize how starvation modulates *Caenorhabditis elegans* vulval cell fate patterning – an environmentally sensitive process, with a nevertheless robust output. Past research has shown many vulval mutants affecting EGF-Ras-MAPK, Delta-Notch and Wnt pathways to be suppressed by environmental factors, such as starvation. Here we aimed to resolve previous, seemingly contradictory, observations on how starvation modulates levels of vulval induction. Using the strong starvation suppression of the Vulvaless phenotype of *lin-3/egf* reduction-of-function mutations as an experimental paradigm, we first tested for a possible involvement of the sensory system in relaying starvation signals to affect vulval induction: mutation of various sensory inputs, DAF-2/Insulin or DAF-7/TGF-*β* signaling did not abolish *lin-3(rf)* starvation suppression. In contrast, nutrient deprivation induced by mutation of the intestinal peptide transporter gene *pept-1* or the TOR pathway component *rsks-1* (the orthologue of mammalian P70S6K) very strongly suppressed *lin-3(rf)* mutant phenotypes. Therefore, physiologically starved animals induced by these mutations tightly recapitulated the effects of external starvation on vulval induction. While both starvation and *pept-1* RNAi were sufficient to increase Ras and Notch pathway activities in vulval cells, the highly penetrant Vulvaless phenotype of a tissue-specific null allele of *lin-3* was not suppressed by either condition. This and additional results indicate that partial *lin-3* expression is required for starvation to affect vulval induction. These results suggest a cross-talk between nutrient deprivation, TOR-S6K and EGF-Ras-MAPK signaling during *C. elegans* vulval induction.

## Introduction

Organismal development is inherently sensitive to environmental variation, and specific environmental conditions, such as nutrient availability or temperature, may also reflect instructive cues controlling growth and other critical developmental decisions, such as developmental timing, diapause entry and production of alternative phenotypes. Developmental integration of environmental cues has become increasingly understood, primarily in the context of nutritional and metabolic regulation of developmental decision, growth and lifespan, such as in *Drosophila* and *C. elegans* (Flatt et al. 2013). In contrast to the understanding of such instructive environmental cues in development, very little is known about how environmental variation impacts other, highly diversified developmental processes, seemingly unaffected by the environment as they maintain their function and corresponding phenotypic outputs when facing such variation. However, such stability of the phenotypic output may go in hand with an underlying flexibility of developmental mechanisms in changing environmental (and genetic) conditions (Félix and Barkoulas 2015; True and Haag 2001; Paaby and Rockman 2014).

*C. elegans* vulval development provides one clear example of a developmental process, which robustly generates an invariant phenotypic output (cell fate pattern) although activities and interactions of underlying molecular signaling pathways are environmentally sensitive (Braendle and Félix 2008; Grimbert and Braendle 2014). *C. elegans* vulval cell fate patterning involves a network of conserved signaling pathways, EGF-Ras-MAPK, Delta-Notch and Wnt pathways that reliably establish a stereotypical cell fate pattern of hypodermal vulval precursor cells (VPCs) (Félix 2012; Sternberg 2005) (Figures 1A and 1B). The VPCs represent a subset of Pn.p cells, P3.p to P8.p,competent to adopt vulval cell fates. Competence of vulval precursor cells is established during L1-L2 stages through expression of the Hox gene *lin-39* and Wnt signals from the posterior (Penigault and Félix 2011; Salser et al. 1993). The key events of the vulval patterning process take place from mid L2 to early L3 stage and involve intercellular signaling between the gonadal anchor cell (AC) and the VPCs (Figures 1A and 1B). In brief, LIN-3/EGF ligand released from the AC induces the primary (1°) vulval cell fate by activating the EGF-Ras-MAPK pathway in P6.p, which receives the highest dose of this signal (Hill and Sternberg 1992; Grimbert et al. 2016). EGF-Ras-MAPK activation induces production of a lateral signal via the Delta-Notch pathway, promoting 2° and inhibiting 1° cell fate in the neighboring cells, P5.p and P7.p (Greenwald et al. 1983; Sternberg and Horvitz 1986; Berset et al. 2001; Yoo et al. 2004). Moreover, a switch from the canonical LET-60/Ras-LIN-45/Raf pathway to a LET-60/Ras-RGL-1-RAL-1/RAL signaling pathway can promote the 2° cell fate in P5.p and P7.p (Zand et al. 2011). The remaining three VPCs, although competent, adopt non-vulval cell fates (3° for P4.p and P8.p, and 3° or 4° for P3.p) as they do not receive sufficient doses of either EGF-Ras-MAPK or Delta-Notch signal. In addition to EGF-Ras-MAPK and Delta-Notch pathways, the canonical Wnt signaling pathway may also be involved in vulval induction: overactivation of the Wnt pathway, e.g. through *pry-1*/*Axin* mutation, increases vulval induction of diverse hypoinduced mutants (Braendle and Félix 2008; Gleason et al. 2002; Seetharaman et al. 2010). *C. elegans* vulval development thus involves a regulatory network of three key molecular cascades and their cross-talk contributes to a reliable and precise patterning output in the presence of both genetic and environmental perturbations (Braendle et al. 2010; Braendle and Félix 2008; Félix and Barkoulas 2012; Gleason et al. 2002; Hoyos et al. 2011; Milloz et al. 2008).

**Figure 1.**
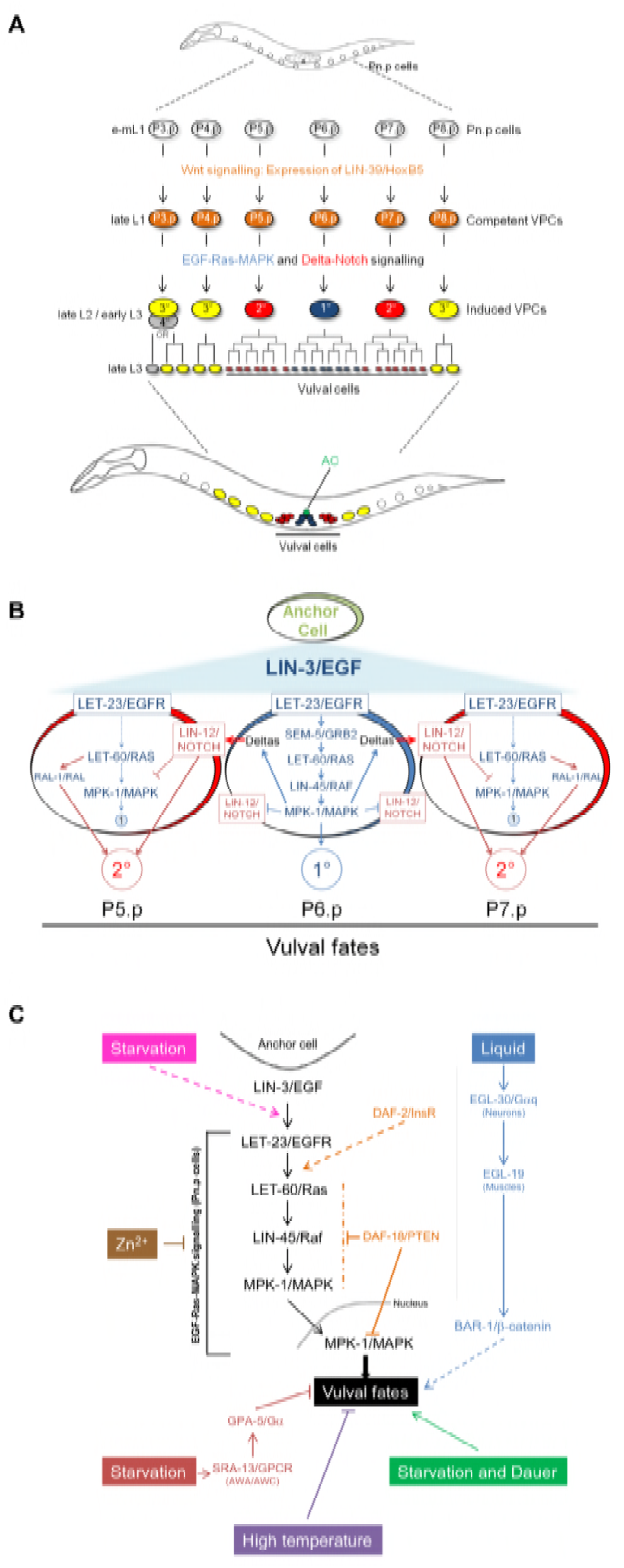
Progression and environmental sensitivity of *C. elegans* vulval cell fate patterning. **(A)** A subset of six Pn.p cells (P3.p to P8.p) acquires competence to form vulval tissue through the expression of LIN-39; however, usually only P5.p, P6.p and P7.p adopt vulval cell fates in a stereotypical 2° −1° −2°. The induction of vulval cell fates is generated by the cross-talk of EGF-Ras-MAPK and Delta-Notch signaling pathways. Distinct vulval cell divisions according to cell fate occur during the L3 stage and generate 22 vulval cells. **(B)** The anchor cell (AC) releases the LIN-3 inductive signal. LIN-3 acts as a morphogen, with P6.p receiving the highest level of LIN-3 causing it to adopt a 1° cell fate (blue). The expression of the Delta ligands in P6.p activates the Delta-Notch pathway in its neighbours, P5.p and P7.p. This activation causes them to adopt a 2° fate (red) and represses the primary fate. A switch from the canonical LET-60/Ras-LIN-45/Raf pathway to a LET-60/Ras-RGL-1-RAL-1 signaling pathway can also promote the 2° cell fate in P5.p and P7.p (not detailed). The competent cells, P4.p and P8.p, adopt a non-vulval 3° fate (not represented), while P3.p may either adopt a 3° fate or a 4° fate, also referred to as F(used) fate. **(C)** Schematic representation of previously reported environmental and physiological inputs affecting *C. elegans* vulval induction (for details, see introduction). Colour coding: Braendle & Félix (2008): pink, Bruinsma et al. (2002): brown, Battu et al. (2003): red, Moghal et al. (2003): blue, Nakdimon et al. (2012): orange, Grimbert & Braendle (2014): purple.

Despite the apparent robustness of *C. elegans* vulval cell fate patterning to environmental perturbations, multiple reports demonstrate that this process is nevertheless responsive to environmental and physiological inputs (Braendle and Felix 2009; Braendle et al. 2008; Félix 2012; Félix and Barkoulas 2012; Sternberg 2005) (summarized in Figure 1C). In early mutagenesis screens, various Vulvaless mutations (*lin-2, lin-3, lin-7, lin-24, lin-33* and *let-23*) were found to be suppressed by starvation and/or dauer passage (Ferguson and Horvitz 1985), suggesting that vulval inductive pathways, such as EGF-Ras-MAPK, are environmentally sensitive. Detailed studies later confirmed that specific chemical elements, such as zinc (Bruinsma et al. 2002; Yoder et al. 2004), and growth conditions indeed modulate vulval inductive signaling. Moghal et al. (2003) reported that liquid culture increases vulval inductive signals via the neuronally expressed heterotrimeric G*α*q protein, EGL-30. The EGL-30/G*α*q signal is transduced via the voltage-gated calcium channel, EGL-19, in muscle cells and its positive effect on vulval induction is mediated by Wnt signaling via BAR-1/β-Catenin (Moghal et al. 2003). Battu et al. (2003) further found evidence for a starvation signal that negatively affects vulval induction via the sensory system. In this study, starvation conditions were found to suppress vulval hyperinduction, for example, induced by *let-60*/*Ras(gf).* This negative starvation effect requires the G-protein-coupled receptor SRA-13 to modulate EGF-Ras-MAPK activity (Battu et al. 2003). Consistent with a sensory-mediated nutritional modulation of vulval signaling, compromised DAF-2/Insulin signaling recapitulates the results that starvation negatively affects vulval induction (Battu et al. 2003; Nakdimon et al. 2012). In addition, another study (Nakdimon et al. 2012) indicates that DAF-18/PTEN negatively regulates EGF-Ras-MAPK activity during vulval induction, further supporting the view that multiple key metabolic and sensory pathways interact with vulval signaling pathways.

Braendle & Félix (2008) examined the effects of diverse environmental conditions (different temperatures, starvation, liquid culture, dauer passage) on vulval induction using a diverse set of known mutations in EGF-Ras-MAPK, Delta-Notch and Wnt pathways. A majority of mutations showed variable penetrance depending on the environment, with starvation and dauer environments showing the most pronounced effects (Braendle and Félix 2008). Consistent with the results of Ferguson & Horvitz (1985), yet seemingly contrary to the results obtained by Battu et al (2003), starvation increased vulval inductive levels, leading to drastic suppression of the Vulvaless phenotypes caused by *lin-3(rf)* and *let-23/egfr(rf)* mutations (Braendle and Félix 2008). Genetic analysis suggests that this starvation signal acts at the level or upstream of LET-23/EGFR (Braendle and Félix 2008). The positive effect of starvation on vulval induction was further confirmed by quantification of pathway activities in starved versus fed wild type animals: starved animals showed a significantly higher EGF-Ras-MAPK pathway activity in the 1° cell, P6.p, whereas Delta-Notch activity was higher in 2° cells, P5.p and P7.p (Braendle and Félix 2008). In addition, the same study found that starvation potentially modulates vulval induction via the Wnt pathway as vulval induction was strongly compromised in starved *bar-1/β-Catenin(0)* animals.

In summary, starvation experienced during larval growth may have both positive (Braendle and Félix 2008; Ferguson and Horvitz 1985) and negative (Battu et al. 2003) effects on vulval induction. While negative starvation effects have been shown to be mediated by the sensory system (Battu et al. 2003), it remains unknown how positive starvation effects are perceived and transduced to modulate vulval induction. In addition, how such, apparently antagonistically acting, starvation signals interact during vulval induction is also unknown.

Addressing these open questions, we have aimed here to quantify and characterize more precisely how starvation affects *C. elegans* vulval cell fate patterning. In particular, we ask how sensory perception of external starvation conditions versus the (internal) animal’s nutrient status affect vulval induction.

## Material and Methods

### Strains and maintenance

Strains were maintained on NGM agar plates (55mm petri dishes, 1.7% agar) carrying a lawn of *E. coli* OP50 (Wood 1988; Brenner 1974). Animals were grown at 20 °C unless indicated otherwise and both mutant and wild type strains were freshly thawed prior to experiments. Our analysis included the *C. elegans* N2 wild type reference strain and the mutants listed below, all of which had been previously isolated and described.

LGI: *egl-30(ad805), che-3(e1124)*

LGII: *let-23(sy1), sra-13(zh13)*

LGIII: *daf-7(e1372), mpk-1(ku1), daf-2(e1370), lin-39(n2110), rsks-1(ok1255), zhIs4 [lip-1::GFP]*

LGIV: *lin-3(e1417), lin-3(n378), lin-3(mf75)*

LGV: *arIs92[egl-17p::NLS-CFP-lacZ, unc-4(+), ttx-3::GFP]*

LGX: *daf-12(rh61rh411), bar-1(mu63), bar-1(ga80), sem-5(n2019), pept-1(Ig601), osm-5(p813)*

The *lin-3(mf75)* allele is a recently described tissue-specific transcriptionally null allele (Barkoulas et al. 2016), where two cis-regulatory elements required for anchor cell expression have been deleted using CRISPR-Cas9 (Dickinson et al., 2013).

### Scoring of vulval phenotypes

The vulval phenotype was observed using Nomarski optics in early to mid L4 individuals, anaesthetized with sodium azide (Wood 1988). We counted the Pn.p progeny and determined their fates as previously described (Braendle and Félix 2008; Sternberg and Horvitz 1986).

### Experimental environments

Experimental populations were age-synchronized by hypochlorite treatment and liquid arrest (27 hours) at the beginning of experiments. Individuals examined in different environments or of different strains were always scored in parallel and derived from populations kept in identical environmental conditions over at least two generations.

For starvation assays, L1 larvae were grown on standard NGM plates until they reached the mid L2 stage (23 hours after L1 transfer back on NGM seeded plates) unless mentioned otherwise. At this stage, animals were washed three times with sterile M9 buffer and transferred on starvation plates, i.e. unseeded NGM plates containing 1mg/ml of ampicillin to prevent bacterial growth. After 48 hours, starved animals were transferred back to regular NGM plates seeded with *E. coli* OP50 and the vulval phenotype was scored when animals had reached the early or mid L4 stage (approximately 15-20 hours later). Control animals were kept on NGM plates seeded with *E. coli* OP50 from L1 to L4. This starvation treatment drastically reduced, yet did not completely stop, developmental progression of worms: after 48 hours most animals had developed into the early to mid-L3 individuals. Note that this starvation treatment did not induce dauer formation.

### RNAi experiments

RNAi by bacterial feeding was performed as described by Timmons et al. (2001). The HT115 bacterial strain carrying the empty RNAi expression vector L4440 served as a negative control. We used RNAi plates composed of standard NGM with 50ug/ml of ampicillin and 1mM of IPTG. Late L2/ early L3 individuals were transferred on RNAi bacteria. The vulval phenotype was scored in L4 individuals of the F1 generation. RNAi clones were selected from Ahringer’s RNAi library (Kamath et al. 2003) or from Vidal’s RNAi library (Rual et al. 2004). The following RNAi clones were used: *pept-1* (clone K04E7.2, Ahringer Library), *let-363* (clone B0261.2a, Ahringer) and *rsks-1* (clone Y47D3A.16, Vidal ORFeome Library).

### Quantification of *pept-1* RNAi effects on Ras and Notch pathway activities

To quantify EGF-Ras-MAPK and Delta-Notch pathway activities in response to *pept-1* RNAi we used previously generated transgenic strains containing integrated transcriptional reporter constructs: the JU480 strain carries the *egl-17::cfp-LacZ* transgene (EGF-Ras-APK activity reporter) derived from the strain GS3582 (Yoo et al. 2004) and the AH142 strain carries the *lip-1::gfp* transgene (Delta-Notch activity reporter) (Berset et al. 2001). Late L2/ early L3 individuals were randomly allocated to *pept-1* RNAi (K04E7.2 clone) or control (L4440 empty vector) plates. EGF-Ras-MAPK and Delta-Notch pathway activities were measured in lethargus L2/early L3 of the F1 generation.

We quantified reporter gene activity in VPCs as previously described (Braendle and Félix 2008; Braendle et al. 2010) (Figure S1). In brief, CFP or GFP signal quantification was performed when individuals had reached the stage of lethargus L2/L3 or early L3. Pn.p cells of live, anesthetized individuals were first identified using DIC imaging, followed by measurement of pixel signal intensity in P5.p, P6.p and P7.p for each individual. Images were acquired using an Olympus BX61 microscope at 40X magnification, equipped with a Coolsnap HQ2 camera. To quantify signal intensity of each cell, we first selected a fixed sub-region of the same size within nuclei of target VPCs and then measured the mean signal intensity of this region. After background subtraction, we used the mean signal intensity as a measure of the corresponding signaling pathway activity in each Pn.p.

### Single Molecule FISH

L3 stage animals were processed to quantify *lin-3/egf* mRNA levels by smFISH as described before (Barkoulas et al. 2013). In brief, they were permeabilized and hybridized with custom-made probes that bind specifically to *lin-3/egf* and *lag-2* mRNAs, labelled with Cy5 and Alexa-594, respectively. The animals were stained with DAPI and placed in GLOX buffer for image acquisition. For each animal, we acquired Z-stacks with a Pixis 1024B camera (Princeton Instruments) mounted onto an upright Zeiss AxioImager M1 with a Lumen 200 metal arc lamp (Prior Scientific). Each Z-stack was composed of 30 images spanning 20.3 µm (0.7 µm in between two acquisitions) on each filter channel. Finally, spot quantification was performed using a custom-made Matlab routine optimized for the anchor cell. Only worms with gonad length > 50 µm were included in the quantifications.

### Statistical Analyses

Data were transformed (e.g. Box-Cox- or log-transformed) when necessary to meet the assumptions of ANOVA procedures (homogeneity of variances and normal distributions of residuals) (Sokal and Rohlf 1981). For post hoc comparisons, Tukey’s honestly significant difference (HSD) procedure was used. Statistical tests were performed using the software programs JMP v13 or SPSS v19.

### Data availability

All strains generated in this study are available upon request.

## Results and Discussion

### Quantifying starvation effects on vulval induction in *lin-3/egf*(*rf*) mutants

Previous studies have used different protocols to examine starvation effects on vulval induction, e.g. animals were exposed to starvation conditions for different periods of time and at different developmental time points (Battu et al. 2003; Braendle and Félix 2008; Ferguson and Horvitz 1985). We therefore first aimed to determine the starvation-sensitive periods of vulval induction using *lin-3(rf)* mutants known to be affected by starvation conditions (Braendle and Félix 2008; Ferguson and Horvitz 1985). Consistent with these earlier studies, a 48h starvation exposure consistently increased vulval induction from late L1/early L2 to early L3 in *lin-3(n378)* animals (Figure 2A). Starvation sensitivity thus extends over the entire period of the vulval cell fate patterning process, with mid-L2 individuals being most sensitive. Next, we asked how the duration of starvation exposure modulates vulval induction. Exposing *lin-3(n378)* animals to different time periods of starvation (0-120h) at the mid L2 stage caused strong suppression of the Vulvaless phenotype when starved for 48 hours or more, low suppression after 19-24h of starvation, and no suppression after a brief (2h) exposure to starvation or early L1 starvation (12h) in liquid medium (Figure 2B). Therefore, the extent of vulval induction depends on the duration of the starvation exposure during specific larval developmental stages.

**Figure 2.**
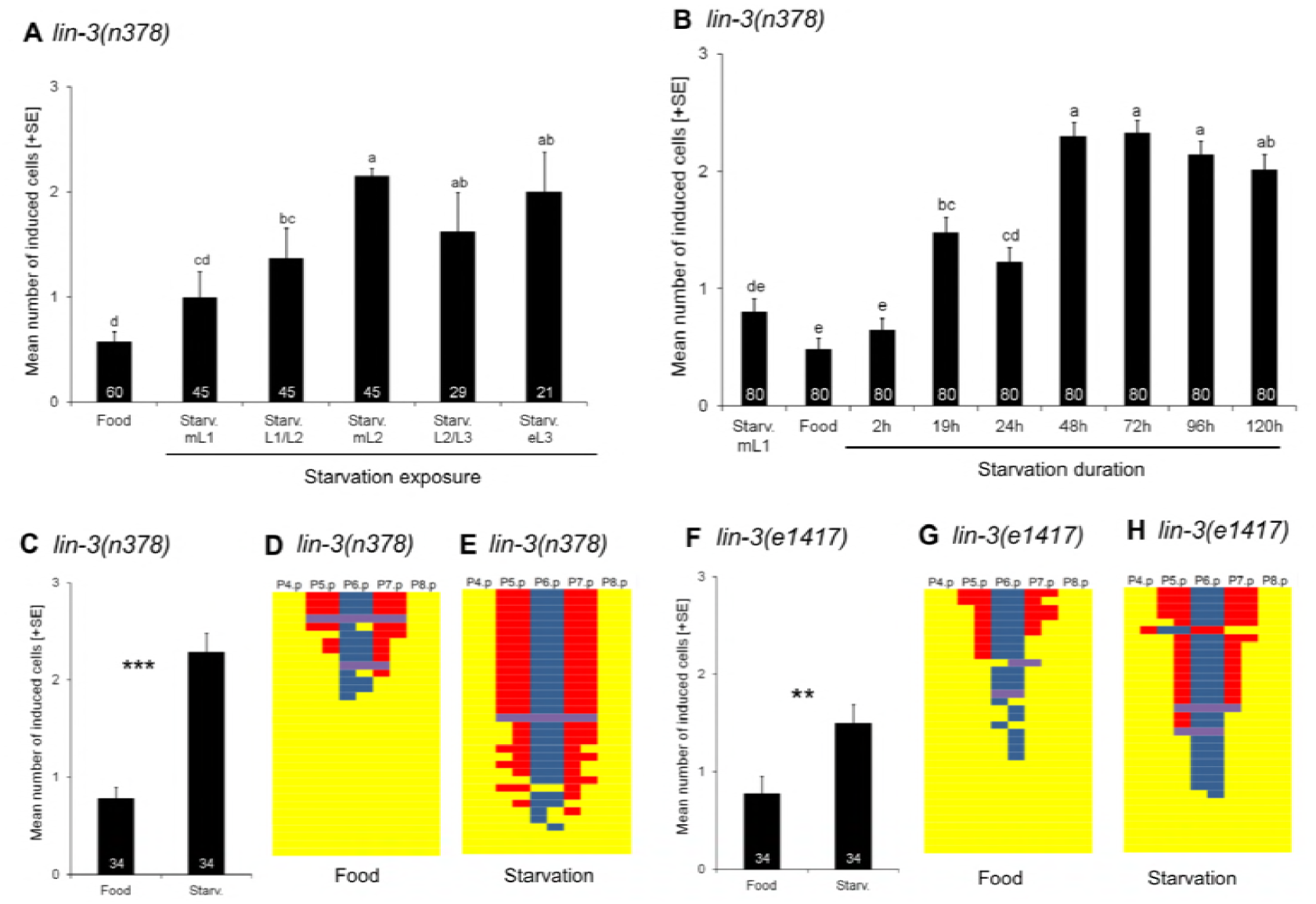
Quantifying starvation effects on vulval induction in *lin-3/egf(rf)* mutants. **(A)** Differences in vulval induction of *lin-3(n378)* animals exposed to starvation (48 hours) at different developmental stages. Starvation suppression of *lin-3(n378)* differed significantly between exposed developmental stages (ANOVA, F_5,239_= 14.02, P < 0.0001), with strongest effects observed from mid L2 to early L3 stages. **(B)** Effects of starvation duration on vulval induction of *lin-3(n378)*. Animals (age-synchronized by egg-laying windows) were exposed to starvation at the mid L2 stage for different time periods except for the first treatment where animals were starved in liquid (M9 buffer) for 12 hours directly after hatching (N=80 per treatment). Starvation duration significantly affected levels of vulval induction (ANOVA, F_5,239_ = 14.02, P < 0.0001) with strongest suppression of *lin-3(n378)* at 48 to 120 hours of starvation. **(C)** Starvation significantly increased vulval induction of *lin-3(n378)* animals relative to control food conditions (ANOVA, F_1,66_ = 33.75, P < 0.0001). **(D, E)** Individual *lin-3(n378)* fate patterns of P4.p to P8.p in **(D)** food versus **(E)** starvation conditions. **(F)** Starvation significantly increased vulval induction of *lin-3(e1417)* animals relative to control food conditions (ANOVA, F_1,66_ = 9.97, P = 0.0024). **(G, H)** Individual *lin-3(e1417)* fate patterns of P4.p to P8.p in **(G)** food versus **(H)** starvation conditions. Values represented by bars **(A,B,C,F)** indicate the mean number of induced vulval cells, also referred to as the vulval index (WT=3 cells induced). Values with different letters indicate significant differences (ANOVA, followed by Tukey’s HSD) (^*^ P < 0.05, ^**^ P < 0.01, ^***^ P < 0.001, ns: non-significant). Numbers displayed in bars represent the number of individuals scored; error bars indicate ± 1 SEM. Vulval cell fate patterns of P4.p to P8.p **(D,E,G,H)** were, whenever feasible, separately inferred for Pn.pa and Pn.pp in cases of half-induced fates. Each line represents the vulval pattern of a single individual, and individuals are ordered from highest to lowest index of vulval induction. Colour coding of vulval cell fates (1° : blue, 2° : red) and non-vulval cell fates (3° : yellow). Induced vulval cells that could not be clearly assigned to either a 1° or 2° fate are shown in purple.

To characterize how starvation of *lin-3(rf)* mutants affects vulval cell lineages, we further scored cell fate patterns in mid-L4 individuals that had been exposed to 48 hours of starvation at the mid-L2 stage. The Vulvaless phenotypes of *lin-3(n378)* and *lin-3(e1417)* were strongly suppressed by starvation (Figures 2C and 2F), similar to previous results (Braendle and Félix 2008). Starvation resulted in an increased mean number of induced vulval cells and a higher proportion of individuals with a canonical 2° −1° −2° cell fate pattern (P5.p to P7.p), and these suppression effects were consistently stronger in *lin-3(n378)* compared to *lin-3(e1417)* (Figures 2C-2H). In *lin-3(n378)*, the proportion of individuals with complete induction increased from 12% in food conditions to 50% after starvation, and most of these individuals showed a correct 2°-1°-2° pattern for P5.p to P7.p (Figures 2D and 2E). Together, these results show that starvation can partly compensate for reduced LIN-3 signaling, confirming the previously reported positive effect of starvation on vulval induction (Braendle and Félix 2008).

### Compromised sensory signaling does not abolish starvation effects on vulval induction

To understand how starvation stimuli are transduced to affect vulval induction, we first tested for an implication of the sensory system in relaying external starvation conditions. Battu et al. (2003) reported that vulval hyperinduction caused by *let-60/ras(gf)* was significantly reduced after starvation of late L1 larvae, indicative of a negative starvation effect on vulval induction. Moreover, this study showed that mutation of the G-protein-coupled receptor *sra-13* or mutations causing sensory signaling defects, such as *osm-5(p813)* and *che-3(e1124)*, abolished starvation sensitivity of *let-60(gf)* animals, implying that observed starvation effects acted on vulval induction via components of the sensory system. We therefore tested whether disruption of the same sensory signaling components also inhibits starvation effects on vulval induction in *lin-3(rf)* mutants. However, we observed that starvation still significantly increased vulval induction of *lin-3(n378)* in *sra-13(zh13), osm-5(p813)* and *che-3(e1124)* backgrounds, comparable to the results of the *lin-3(n378)* single mutant (Table 1). Therefore, in contrast to the results of Battu et al. (2003), these sensory mutants did not abolish starvation suppression of the Vulvaless phenotype in *lin-3(n378)* mutants.

**Table 1.** Effects of starvation treatment on vulval induction level of different mutants of the sensory system. Statistical tests (One-Way-ANOVAs) were performed for each mutant separately testing for the effect of treatment (food vs starvation) on the number of vulval cells induced (^*^ P < 0.05, ^**^ P < 0.01, ^***^ P < 0.001, ns: non-significant). To facilitate comparison, measurements for *lin-3(e1417)* and *lin-3(n378)* are also included (i.e. same data as in Figures 2C and 2F). *egl-17::cfp* *lip-1::gfp*

| Genotype | Treatment | Induced cells<br>(Mean ± SE) | N |
| --- | --- | --- | --- |
| *lin-3(e1417)* | Food | 0.77 ± 0.12 | 34 |
| | Starvation | 1.50 ± 0.18 *** | 34 |
| *lin-3(n378)* | Food | 0.78 ± 0.12 | 34 |
| | Starvation | 2.28 ± 0.20 *** | 34 |
| *che-3(e1124); lin-3(n378)* | Food | 0.94 ± 0.04 | 60 |
| | Starvation | 2.53 ± 0.21 *** | 37 |
| *lin-3(n378); osm-5(p813)* | Food | 0.91 ± 0.13 | 60 |
| | Starvation | 2.58 ± 0.13 *** | 30 |
| *sra-13(zh13); lin-3(n378)* | Food | 0.86 ± 0.13 | 60 |
| | Starvation | 2.07 ± 0.15 *** | 43 |
| *daf-7(e1372); lin-3(e1417)* | Food | 0.26 ± 0.11 | 32 |
| | Starvation | 0.92 ± 0.20 ** | 35 |
| *daf-2(e1370); lin-3(n378)* | Food | 0.01 ± 0.01 | 60 |
| | Starvation | 1.13 ± 0.18 ** | 60 |
| *lin-3(n378); daf-12(rh61rh411)* | Food | 0.34 ± 0.09 | 60 |
| | Starvation | 1.77 ± 0.31 *** | 16 |

We next tested for an implication of TGF-β and Insulin signaling: two key signaling pathways involved in sensory transduction of environmental stimuli, which mediate diverse developmental and metabolic responses to changes in nutritional status, e.g. during dauer formation (Fielenbach and Antebi 2008) or germline proliferation (Dalfo et al. 2012; Michaelson et al. 2010). The activity of the TGF-β and insulin pathways are reduced upon starvation. Reducing TGF-β pathway activity did not abolish starvation effects on vulval induction as shown by the double mutant *daf-7(e1372); lin-3(e1417)* (Table 1). Similarly, reduced activity of the DAF-2/Insulin receptor caused by the *daf-2(e1370)* mutation did not abolish starvation effects on *lin-3(n378)* mutants (Table 1). However, vulval induction in *daf-2(e1370) lin-3(n378)* double mutants was significantly lower than in the single *lin-3(n378)* mutant, both in food and starvation conditions. This result is consistent with previous observations that reduced Insulin signaling may lower vulval induction in sensitized backgrounds, such as *let-60/Ras(gf)* (Battu et al. 2003; Nakdimon et al. 2012). To confirm that TGF-β and Insulin signaling do not mediate positive starvation effects on vulval induction, we further examined the role of a central genetic component, the DAF-12 steroid receptor, integrating downstream effects of these two signals (Fielenbach and Antebi 2008) using the Daf-d mutant, *daf-12(rh61rh411)* (Antebi et al. 2000). Again, as for *daf-2* and *daf-7,* this mutant did not abolish positive starvation effects on *lin-3(n378)* vulval induction (Table 1).

Taken together, these results indicate that Insulin and TGF-β pathways, and thus by implication, environmental signal transduction via these pathways, do not play major roles in mediating positive starvation effects on vulval induction. However, we confirm previous observations (Battu et al. 2003; Nakdimon et al. 2012) that reduced DAF-2 Insulin activity may lower vulval induction in sensitized backgrounds, including *lin-3(rf)* mutants, in both food and starvation conditions. Yet, in our experiments, starvation nevertheless increased vulval induction of *daf-2(e1370); lin-3(n378*) animals, suggesting that the negative starvation effect (Battu et al. 2003; Nakdimon et al. 2012) on vulval induction is significantly weaker than the positive starvation signal. Therefore, while Insulin signaling and sensory perception seem to mediate a mild negative starvation signal (Battu et al. 2003; Nakdimon et al. 2012), we here provide evidence for an additional positive starvation signal acting independently of Insulin and sensory signaling. Moreover, starvation thus appears to induce multiple antagonistic effects on *C. elegans* vulval induction.

### Nutrient-deprivation caused by *rsks-1* and *pept-1* mutations strongly increase vulval induction in *lin-3(rf)*

To test whether observed starvation effects on vulval induction might be mediated by internal perception of the organism’s nutritional status, we focused on a conserved key regulator of cell growth and proliferation via nutrient sensing, the LET-363/TOR (target of rapamycin) pathway (Hietakangas and Cohen 2009; Jia et al. 2004). RNAi knock-down of *let-363* and *rsks-1/S6K* (encoding the *C. elegans* orthologue of mammalian P70S6K, a known TORC1 target) (Laplante and Sabatini 2012), significantly increased vulval induction of *lin-3(n378)* animals in food conditions (Figure 3A). Moreover, RNAi knockdown of *pept-1* (encoding an intestinal oligopeptide transporter) (Fei et al. 1998), causing nutrient-deprived animals through amino acid starvation (Meissner et al. 2004), increased vulval induction of *lin-3(n378)* even more strongly (Figure 3A).

**Figure 3.**
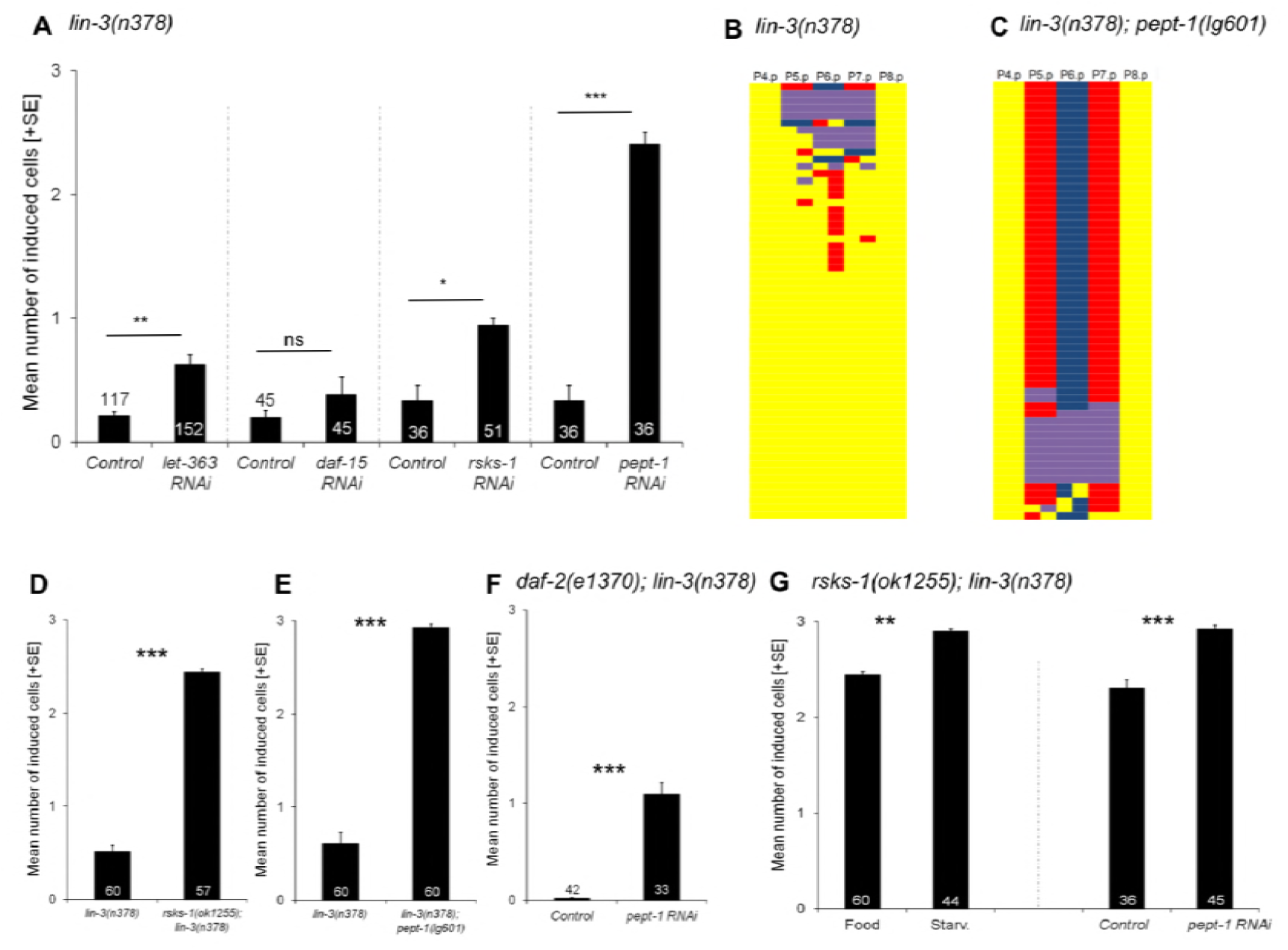
Nutrient-deprivation caused by *rsks-1* and *pept-1* mutations strongly increases vulval induction in *lin-3/egf(rf).* (A) Effects of *let-363, daf-15, rsks-1* and *pept-1* RNAi versus control RNAi treatment (empty vector strain, *E. coli* HT115) of *lin-3(n378)* animals (food conditions). RNAi knock-down of *let-363, rsks-1* and *pept-1* (but not *daf-15*) significantly increased vulval induction (ANOVA, all P < 0.01). **(B)** *rsks-1(ok1255)* strongly increased vulval induction of *lin-3(n378)* animals (ANOVA, F_1,115_ = 48.42, P < 0.0001). **(C)** *pept-1(lg601)* strongly increased vulval induction of *lin-3(n378)* animals (ANOVA, F_1,118_ = 335.58, P < 0.0001). **(D, E)** Individual fate patterns of P4.p to P8.p in **(D)** *lin-3(n378)* versus **(E)** *lin-3(n378); pept-1(lg601)* individuals. Vulval cell fate patterns of P4.p to P8.p **(D,E)** were, whenever feasible, separately inferred for Pn.pa and Pn.pp in cases of half-induced fates. Each line represents the vulval pattern of a single individual, and individuals are ordered from highest to lowest index of vulval induction. Colour coding of vulval cell fates (1° : blue, 2° : red) and non-vulval cell fates (3° : yellow). Induced vulval cells that could not be clearly assigned to either a 1° or 2° fate are coded in purple. **(F)** Effects of *pept-1* RNAi versus control RNAi (empty vector strain, *E. coli* HT115) in *daf-2(e1370); lin-3(n378)* (food conditions). RNAi knock-down of *pept-1* significantly increased vulval induction (ANOVA, F_1,73_ = 133.57, P < 0.0001). **(G) 1)** Starvation significantly increased vulval induction of *rsks-1(ok1255); lin-3(n378)* animals relative to control food conditions (ANOVA, F_1,102_= 11.62, P = 0.0009) **2)** Effects of *pept-1* RNAi versus control RNAi (empty vector strain, *E. coli* HT115) in *rsks-1(ok1255); lin-3(n378)* (food conditions). RNAi knock-down of *pept-1* significantly increased vulval induction (ANOVA, F_1,79_ = 25.93, P < 0.0001).

These RNAi effects thus mimicked starvation effects on *lin-3(rf)*, consistent with the idea that TOR pathway components and targets mediate starvation effects on vulval induction. We further tested whether RNAi knock-down of an additional TOR pathway member *daf-15* (RAPTOR) affects vulval induction using the *lin-3(n378)* mutant. RNAi of *daf-15* did not cause any suppression of the Vulvaless phenotype of *lin-3(n378)* (Figure 3A).

To consolidate the above results, we constructed double mutants of *lin-3(n378)* with *rsks-1(ok1255)* and *pept-1(lg601)*, respectively. The two double mutants exhibited extremely strong suppression of the *lin-3(n378*) Vulvaless phenotype (Figures 3B to 3E). The suppression was particularly strong in *lin-3(n378); pept-1(lg601)* where mean vulval induction approached wild type levels (2.93 ± 0.03 cells induced) (Figure 3E), and 70% of individuals generated a correct vulval patterning sequence of 2°-1°-2° cell fates for P5.p to P7.p compared to <5% of *lin-3(n378)* individuals (Figures 3B and 3C). Similarly, we also observed strong suppression of the Vulvaless phenotype for another *lin-3* allele, *e1417,* when placed in a *pept-1(lg601)* background [*lin-3(e1417); pept-1(lg601)*: 2.13±0.17 cells induced, N=34; *lin-3(e1417)*: 0.94 ± 0.13 cells induced, N=45; ANOVA, F_1,77_ = 30.09, P < 0.0001]. Of note, however, neither *pept-1(lg601)* or *rsks-1(ok1255)* single mutants exhibited vulval hyperinduction or other vulval defects (N>300/strain). As compromised PEPT-1 activity leads to strongly nutrient-deprived animals, even in a food-rich environment (Meissner et al. 2004; Geillinger et al. 2014), these results are further consistent with our observation that sensory perception of external starvation conditions is not required to increase vulval induction of *lin-3(rf)* animals (Table 1).

Given that reduced amino acid availability caused by *pept-1(lg601)* primarily acts through TOR signaling (Meissner et al. 2004; Korta et al. 2012) and that RNAi of *rsks-1* similarly mimics starvation effects, we conclude that the observed positive starvation modulation of vulval induction likely involves TOR-S6K signaling.

Although TOR and DAF-2 Insulin signaling may largely act independently of each other, cross-talk between the two pathways has also been reported in some biological processes (Korta et al. 2012; Spanier et al. 2010; Meissner et al. 2004). We therefore tested how *pept-1* RNAi affects vulval induction in *daf-2(e1370); lin-3(n378)* animals: *pept-1* RNAi increased vulval induction in a *daf-2(e1370); lin-3(n378)* double mutants, but to a lesser extent than in *lin-3(n378)* single mutants (Figure 3F). This result also tightly mimics the observed starvation effects on vulval induction in *daf-2(e1370); lin-3(n378)* animals (Table 1), corroborating the interpretation that compromised DAF-2 signaling causes a slight reduction of vulval induction, as indicated by earlier studies (Battu et al. 2003; Nakdimon et al. 2012). Increased vulval induction caused by *pept-1* RNAi or starvation thus largely acts independently of DAF-2/IIR signaling.

In both RNAi and mutant experiments, disruption of PEPT-1 activity caused a stronger increase in *lin-3(n378)* vulval induction compared to disruption of RSKS-1 activity (Figures 3B and 3C). Therefore, nutrient deprivation caused by reduced PEPT-1 activity did not precisely recapitulate the effects of the *rsks-1* mutant, which could indicate that factors other than *rsks-1* mediate effects of nutrient deprivation on vulval induction. Consistent with this idea, we found that *rsks-1(ok1255); lin-3(n378)* double mutants exposed to either *pept-1* RNAi or starvation exhibited a slight, but significant increase in vulval induction compared to control animals (Figure 3G). Positive effects of nutrient deprivation on vulval induction may thus involve additional factors other than TOR-S6K signaling.

Overall, nutrient deprivation – caused by external starvation through absence of food or by internal, physiological starvation due to *pept-1* mutation or RNAi – have very similar effects, generating an overall net increase in vulval induction.

### *pept-1* RNAi increases Ras and Notch pathway activities in vulval precursor cells

To address how physiological starvation caused by *pept-1* RNAi affects the vulval signaling network, we focused on the main inductive pathway, the Ras pathway. Starvation conditions have previously been shown to increase EGF-Ras-MAPK activity in the 1° cell, P6.p, and Delta-Notch activity in the 2° cells, P5.p and P7.p of the wild type strain N2 (Braendle and Félix 2008). We therefore tested whether *pept-1* RNAi mimicked these starvation effects using the same fluorescent reporters, *egl-17::cfp* to quantify Ras pathway activity (Yoo et al. 2004) and *lip-1::gfp* to quantify Delta-Notch activity (Berset et al. 2001). *pept-1* RNAi effects closely mirrored previously quantified starvation effects on report gene activities (Braendle and Félix 2008), with *pept-1* RNAi increasing Ras pathway activity in P6.p and increasing Notch pathway activity in P5.p (but not P7.p) (Figures 4A and 4B; Table 2). Consistent with these changes, we also observed that *pept-1* RNAi reduced Delta-Notch activity in P6.p while reducing Ras activity in P5.p and P7.p (Figures 4A and 4B; Table 2). *pept-1* RNAi thus tightly recapitulated previously observed starvation effects on activities of the two key inductive vulval signaling pathways (Braendle and Félix 2008). Observed changes in pathway activities may be explained by distinct effects on Ras and Notch pathways, or by effects mediated solely by the Ras pathway, given the cross-talk between the two pathways (Yoo et al. 2004).

**Figure 4.**
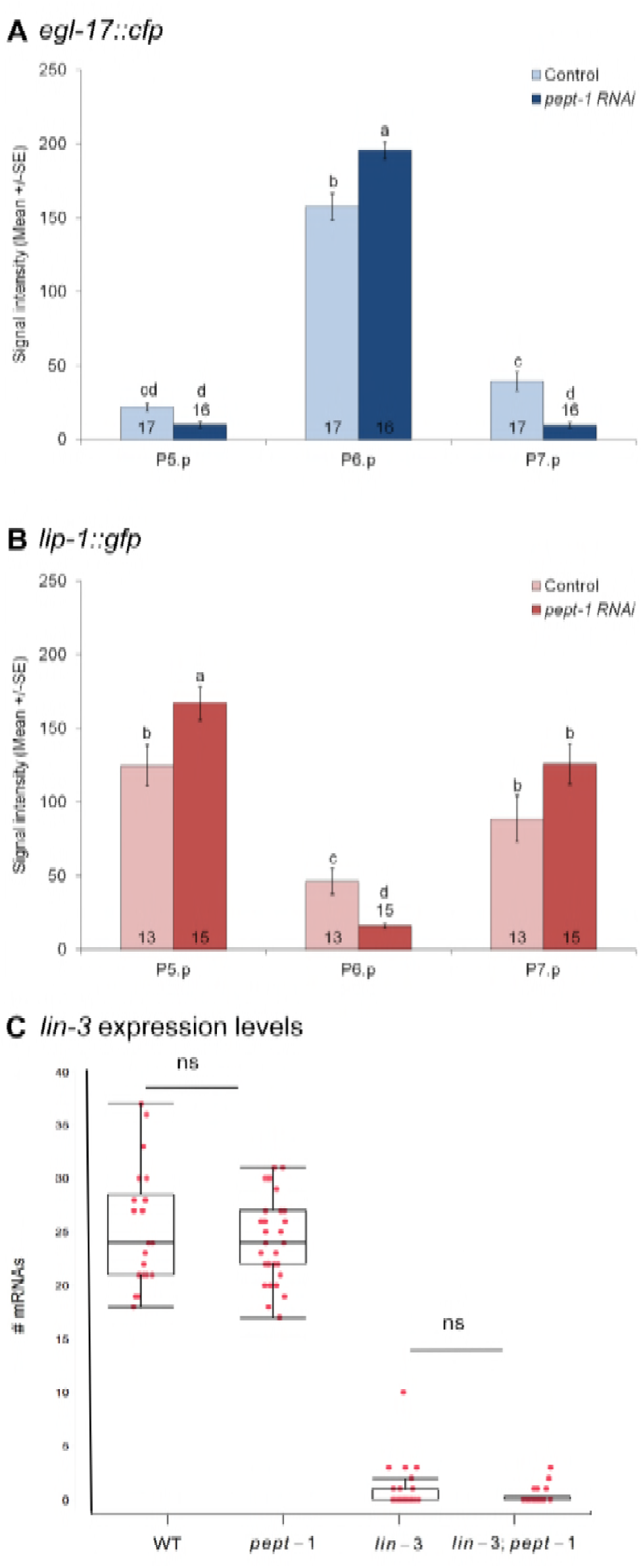
Reduced activity of PEPT-1 increases Ras and Notch pathway activities but does not alter *lin-3* transcript levels. **(A,B)** Effects of *pept-1* RNAi on the level of the transcriptional reporters *egl-17::cfp* (EGF-Ras-MAPK activity) (Yoo et al. 2004) and *lip-1::gfp* (Delta-Notch activity) (Berset et al. 2001) quantified in lethargus L2/L3 and early L3 stages. **(A)** Mean signal (pixel) intensity of the EGF-Ras-MAPK pathway reporter, *egl-17::cfp* in P5.p to P7.p. Values indicate Least Square Means for the interaction *cell* x *RNAi treatment* (F_2,62_ = 15.52, P < 0.0001) (Table 2A). **(B)** Mean signal (pixel) intensity of the Delta-Notch pathway reporter, *lip-1::gfp* in P5.p to P7.p. Values indicate Least Square Means for the interaction *cell* x *RNAi treatment* (F_2,52_ =13.73, P < 0.0001) (Table 2B). For complete statistical analysis and results, see Table 2. Values with different letters indicate significant differences (Tukey’s HSD). Numbers displayed in bars represent the number of individuals scored; error bars indicate ± 1 SEM. **(C)** Boxplot showing *lin-3* mRNA levels in different genetic backgrounds [wild type, *pept-1(lg601), lin-3(e1417)* and *lin-3(e1417); pept-1(lg601)*] is displayed. The *pept-1* mutation does not alter the *lin-3* expression level (mean = 24.38±0.67; n=34) compared to the wild type (mean = 25.36±0.35; n=22) (ANOVA, F_1,54_ = 0.49, P = 0.49). As expected, there is little expression in *lin-3(e1417)* mutant animals (Mean: 0.77±0.35, n=31) but this low expression is not rescued in *lin-3; pept-1* double mutant animals (Mean: 0.36±0.17; n=22) (ANOVA, F_1,751_= 0.46, P = 0.50).

**Table 2.** Effects of *pept-1* RNAi on EGF-Ras-MAPK reporter *egl-17::cfp*. **(A) and Delta-Notch reporter *lip-1::gfp* (B).** Results of statistical tests for transcriptional reporter assays (Figure 5). We used an ANOVA testing for the fixed effects of *RNAi treatment, Individual* (nested in *RNAi treatment*), *Pn.p cell* and the interaction between *RNAi treatment* and *Pn.p cell* using mean signal intensity as a response variable. Including the effect *Individual* (*RNAi treatment*), allowed controlling for the non-independence of signal intensity in P5.p, P6.p, and P7.p measured in a given individual. Data was log-transformed for analysis.

| Source | DF | Sum of Squares | F Ratio | P |
| --- | --- | --- | --- | --- |
| *RNAi Treatment* | 1 | 26.56 | 0.04 | 0.8413 |
| *Individual (RNAi Treatment)* | 31 | 2792.82 | 0.14 | 1.0000 |
| *Pn.p cell* | 2 | 538247.46 | 409.63 | < 0.0001 |
| *Pn.p cell x RNAi Treatment* | 2 | 20392.73 | 15.52 | < 0.0001 |
| Error | 62 | 40733.51 |  |  |

**B) *lip-1::gfp***
| Source | DF | Sum of Squares | F Ratio | P |
| --- | --- | --- | --- | --- |
| *RNAi Treatment* | 1 | 456.59 | 0.34 | 0.5607 |
| *Individual (RNAi Treatment)* | 26 | 24120.94 | 0.70 | 0.8403 |
| *Pn.p cell* | 2 | 200764.78 | 75.39 | < 0.0001 |
| *Pn.p cell x RNAi Treatment* | 2 | 36553.06 | 13.73 | < 0.0001 |
| Error | 52 | 69237.88 |  |  |

### Starvation sensitivity of vulval induction requires minimal *lin-3/egf* expression

To further characterize how physiological starvation interacts with vulval induction to modulate inductive levels, we characterized effects of *pept-1* RNAi on several components of the core EGF-Ras-MAPK cascade using various reduction-of-function mutants (Table 3). In addition to *lin-3(rf)* mutations, *pept-1* RNAi suppressed, to different degrees, the Vulvaless phenotypes of *let-23(sy1), sem-5(n2019)* and *mpk-1(ku1)* (Table 3). Although these results suggest that nutrient deprivation due to *pept-1* RNAi may act downstream of MAPK, interpretation of these effects is limited given that these mutations do not represent null alleles, most of which are homozygous lethals. We therefore used a double mutant *let-23(sy1); lin-3(n378)* that exhibits a fully penetrant Vulvaless phenotype, indicating absence or very low levels of basal Ras activity (Braendle and Félix 2008). Vulval induction of *let-23(sy1); lin-3(n378)* animals treated with *pept-1* RNAi remained virtually unaltered compared to controls (Table 3), with only 4/60 individuals showing some induced vulval cells (versus 0/45 individuals in food conditions). Very similarly, a recently constructed tissue-specific null allele of *lin-3/EGF, mf75* (Barkoulas et al. 2016), did not show any sensitivity to *pept-1* RNAi, suggesting that loss of basal EGF-Ras-MAPK activity due to complete lack of LIN-3 abolishes starvation effects. These results indicate that nutrient deprivation via PEPT-1 modulates vulval induction upstream of LIN-3 or act in parallel to reinforce expression of downstream target genes of the Ras pathway (as seen with transcriptional reporters).

**Table 3.**
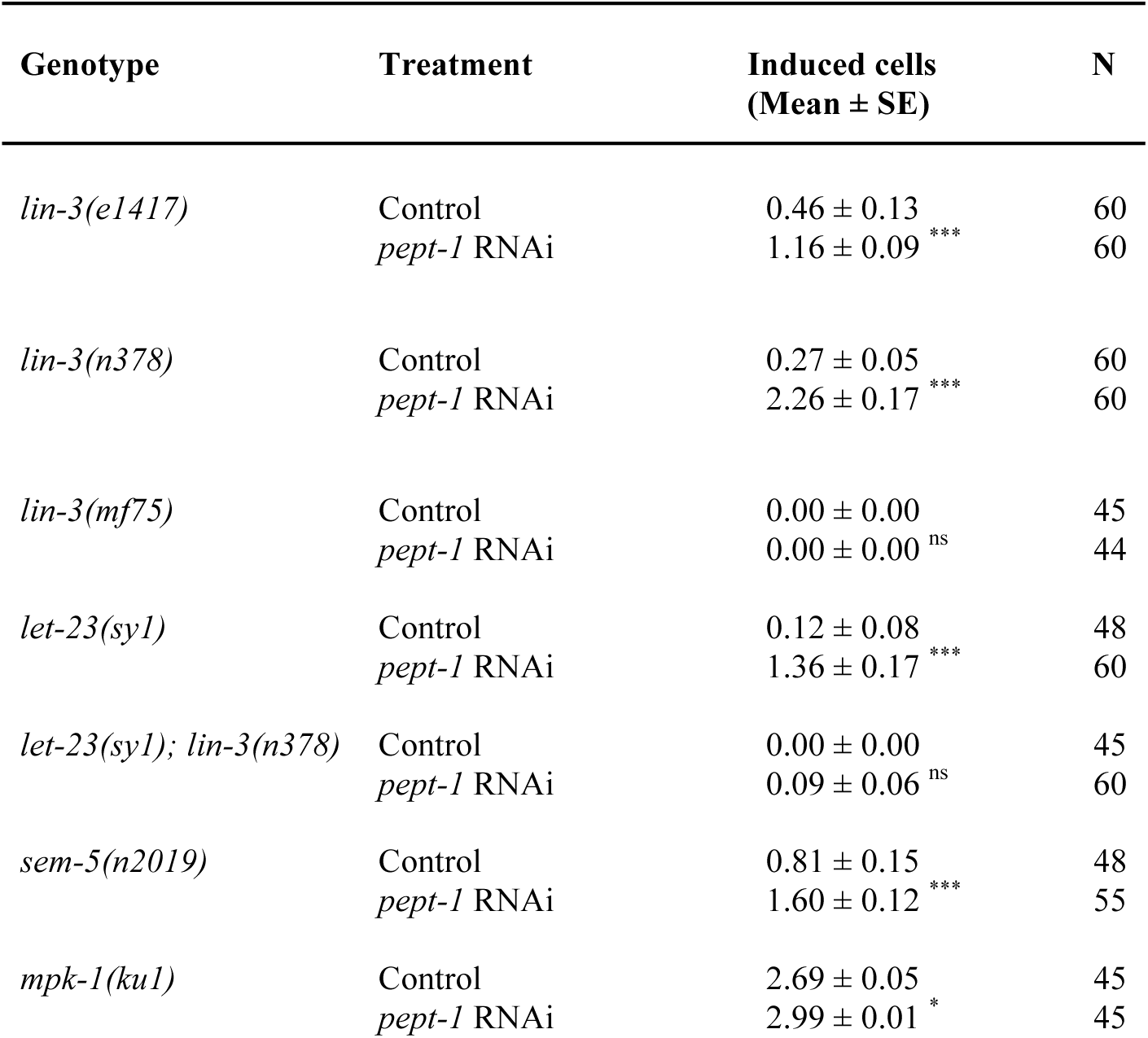
Effects of *pept-1* RNAi treatment on vulval induction level of different mutants of the EGF-Ras-MAPK signaling cascade. Statistical tests (One-Way-ANOVAs) were performed for each mutant separately testing for the effect of treatment (control versus *pept-1* RNAi) on the number of vulval cells induced (^*^ P < 0.05, ^**^ P < 0.01, ^***^ P < 0.001, ns: non-significant).

Given the above results, we next tested whether nutrient deprivation caused by the *pept-1* mutation may alter levels of *lin-3* expression in the anchor cell. Using single molecule FISH, we therefore compared *lin-3* transcript number in wild type, *pept-1(lg601), lin-3(e1417)* and *lin-3(e1417); pept-1(lg601)* animals (Figure 4C). However, there was no effect of the *pept-1* mutation on *lin-3* transcript numbers in either a wild type genetic background or in a background with low transcriptional level of *lin-3*. Therefore, increased vulval induction of *lin-3(e1417)* (Figure 4C) caused by *pept-1* mutation is not due to increased transcription of *lin-3* in the anchor cell. This result, together with the observation that vulval induction of *lin-3(mf75),* a genetic background with no expression of *lin-3* in the anchor cell, is insensitive to *pept-1* RNAi, suggests that starvation effects on vulval induction can only manifest under minimal EGF-Ras-MAPK activity, corresponding to the situation where one or more *lin-3* transcripts are present in the AC. One possible scenario is that the effect of such low doses of *lin-3* on vulval induction can then become amplified under nutrient deprivation, e.g. at the level of the LIN-3 protein and its secretion, or at the level or downstream of the EGF receptor (LET-23)

### Role of BAR-1 activity in starvation modulation of vulval induction

In addition to the EGF-Ras-MAPK, starvation effects on vulval induction may also act through additional signaling pathways, in particular, the Wnt pathway. Previous results suggest that partial redundancy between Ras and Wnt pathway in vulval induction may allow for compensatory interplay when one of the two pathways is perturbed due to mutation (Gleason et al. 2002) or exposure to specific environments, such as liquid (Moghal et al. 2003) or starvation (Braendle and Félix 2008). Gleason et al. (2002) observed that Wnt pathway overactivation (by mutation of *pry-1*) may partly compensate for reduced EGF-Ras-MAPK activity during vulval induction, e.g. in *let-23(rf)* mutants. Braendle & Félix (2008) further found that the vulval hypoinduction phenotype of the null mutant of *bar-1*/*β-Catenin* (allele *ga80*) (Eisenmann et al. 1998) was strongly aggravated under starvation conditions. Moreover, the highly penetrant Vulvaless phenotype of *lin-3(n378); bar-1(ga80)* double mutants was not suppressed by starvation, implying that Wnt pathway activity acting via *bar-1*/*β-Catenin* is required for starvation suppression of *lin-3(rf)*. However, it remained unclear to what extent starvation compromised induction versus competence of vulval cells in *bar-1(ga80)* because in the study of Braendle & Félix (2008) animals were exposed to starvation from the late L1 stage – a stage during which competence of vulval precursor cells is acquired, yet induction does not yet take place. To distinguish these two scenarios, we exposed *bar-1(ga80)* animals to starvation in late L1 versus mid-L2 stage and found that mid-L2 starvation did not alter vulval induction whereas late L1 starvation caused strong hypoinduction, primarily due to Pn.p fusion, i.e. loss of competence of vulval precursor cells, including P5.p to P7.p (Figures 5A-E). Late L1 starvation therefore aggravated *bar-1(ga80)* fusion defects prior to vulval induction, so that this starvation treatment cannot be used to evaluate starvation effects on vulval inductive levels. In contrast, starvation at a later time point, at mid-L2 – covering the time period during which vulval induction is most sensitive to starvation – slightly, although not significantly, increased vulval induction of *bar-1(ga80)* animals (Figure 3A). Yet, *pept-1* RNAi treatment of *bar-1(ga80)* clearly and significantly increased vulval induction (Figure 5F), indicating that such a prolonged starvation stimulus can indeed increase vulval induction in the absence of BAR-1 activity. However, when we exposed *lin-3(n378); bar-1(ga80)* double mutants to *pept-1* RNAi treatment (Control: 0.00±0.00 cells induced, N=44; *pept-1* RNAi: 0.03 ± 0.01 cells induced, N=59), or 48 hours of starvation at the mid-L2 stage (Food: 0.00±0.00 cells induced, N=45; Starvation: 0.01 ± 0.01 cells induced, N=47), we observed no starvation suppression of this highly penetrant Vulvaless phenotype. This result suggests, similar to the results obtained for other highly penetrant *Vulvaless* mutants [e.g. *let-23(sy1); lin-3(n378)* or *lin-3(mf75)*], that a very strong reduction in inductive signals (including contribution from BAR-1/Wnt) cannot be rescued by starvation effects. Nevertheless, given that vulval induction of the *bar-1* null allele *ga80* increases in response to starvation, we conclude that Wnt pathway/BAR-1 activity is not essential to mediate positive starvation effects on vulval inductive levels.

**Figure 5.**
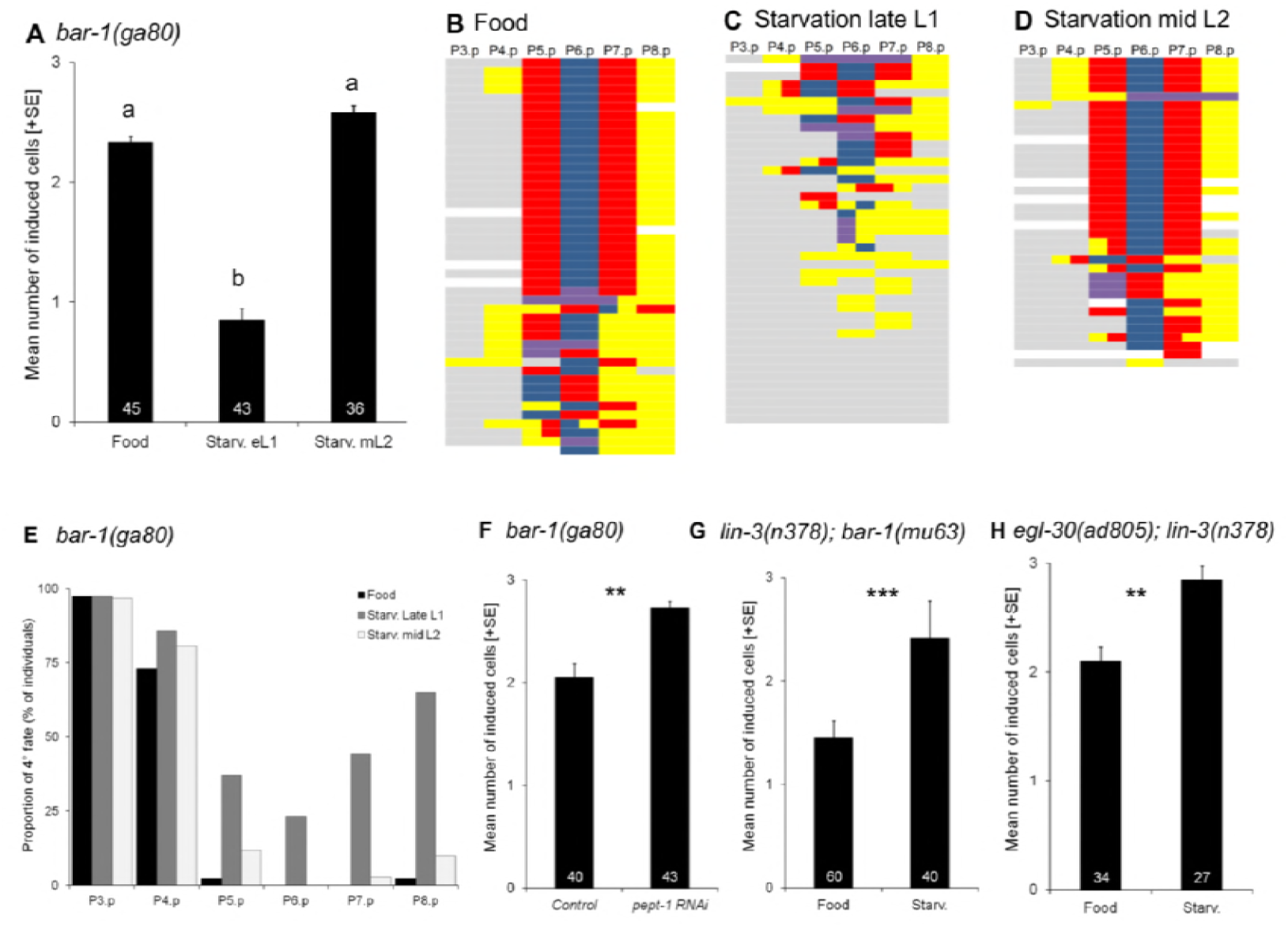
Role of BAR-1 activity in starvation modulation of vulval induction. **(A)** Differences in vulval induction of *bar-1(ga80)* animals exposed to 48 hours of starvation in late L1 versus mid L2 (and versus control food conditions). Starvation at the late L1 stage, but not at the mid L2 stage, significantly reduced vulval induction of *bar-1(ga80)* animals (ANOVA, F_2,121_ = 50.27, P < 0.0001). **(B-D)** Individual *bar-1(ga80)* fate patterns of P3.p to P8.p in **(B)** food versus starvation conditions: late L1**(C)**or mid L2 starvation **(D)**. Vulval cell fate patterns of P3.p to P8.p **(B-D)** were, whenever feasible, separately inferred for Pn.pa and Pn.pp in cases of half-induced fates. Each line represents the vulval pattern of a single individual, and individuals are ordered from highest to lowest index of vulval induction. Colour coding of vulval cell fates (1° : blue, 2° : red) and non-vulval cell fates (3° : yellow, 4° : grey). Induced vulval cells that could not be clearly assigned to either a 1° or 2° fate are coded in purple; non-induced cells that could not be clearly assigned a 3° or° fate are coded in white. **(E)** Proportion of Pn.p cells with a 4° (Fused) fate in control, starvation-exposed late L1, starvation-exposed mid L2 *bar-1(ga80)* individuals (from the same experiment as shown in **(A)**). The proportions of 4° fates for each of P5.p to P7.p cell were higher after L1 starvation (P5.p: 37%, P6.p: 23%, P7.p: 44%) compared to control (P5.p: 2%, P6.p and P7.p: 0%) and L2 starvation (P5.p: 12%, P6.p: 0%, P7.p: 3%). **(F)** *pept-1* RNAi increases vulval induction of *bar-1(ga80)* animals (ANOVA, F_1,81_ = 14.47, P = 0.0003). **(G)** Starvation increased vulval induction of *lin-3(n378); bar-1(mu63)* relative to control food conditions (ANOVA, F_1,98_ = 18.84, P < 0.0001). **(H)** Starvation increased vulval induction of *egl-30(ad805); lin-3(n378)* relative to control food conditions (ANOVA, F_1,59_ = 7.28, P = 0.0091).

A previous study has further shown that liquid culture of animals suppressed the Vulvaless phenotype of *lin-3(rf)* and other EGF-Ras-MAPK mutations through the Wnt pathway (Moghal et al. 2003) (Figure 1C). This environmental effect on vulval induction is based on activation of the heterotrimeric Gαq protein, EGL-30, which acts with muscle-expressed EGL-19 to promote vulval induction downstream or in parallel to LET-60/Ras in a Wnt-dependent manner (Moghal et al. 2003). The liquid effect on vulval induction is abolished when the Wnt pathway is mildly compromised (Moghal et al. 2003), i.e. in a *bar-1(mu63)* context (Maloof et al. 1999). We therefore asked whether starvation effects on *lin-3(rf)* could be mediated in the same fashion. However, starvation did not alter vulval induction of *lin-3(n378)* in a *bar-1(mu63)* background (Figure 5G). Furthermore, loss of *egl-30/Gαq* (Brundage et al. 1996) did not abolish starvation effects on vulval induction of *lin-3(n378)* (Figure 5H). Therefore, the positive starvation effects on vulval induction seem to be distinct from previously reported effects of liquid culture promoting vulval induction (Moghal et al. 2003).

## Conclusions

Analysis of starvation sensitivity of *C. elegans* vulval induction using mutant analysis and quantification of pathway activities allowed us to clarify how a specific environmental condition modulates *C. elegans* vulval induction. We present quantitative analyses of starvation effects on vulval induction using *lin-3(rf)* mutants, demonstrating that starvation exposure of >19h exerts a strong positive effect during the entire period of vulval induction, spanning from mid-L2 to early L3 stages. By testing various candidate mechanisms that could transduce and elicit observed starvation effects, we find that compromised sensory signaling through mutation of *daf-2* (Insulin) or *daf-7* (TGF-*β*) does not abolish *lin-3(rf)* starvation suppression. Instead, nutrient-deprived animals induced by mutation of the intestinal peptide transporter *pept-1* or by the TOR pathway component *rsks-1* (in a food-rich environment) strongly mimicked suppression of *lin-3(rf)* by external starvation. Moreover, we find that *pept-1* RNAi is sufficient to increase both EGF-Ras-MAPK and Delta-Notch pathway activities in Pn.p cells. These results provide evidence for a link between nutrient deprivation, TOR-S6K and EGF-Ras-MAPK signaling during *C. elegans* vulval induction. It remains unclear how nutrient deprivation feeds into genetic pathways regulating vulval development, and we do not know whether this phenomenon represents a specific mechanism to fine-tune levels of vulval induction in response to the environment. In any case, the observed starvation modulation of *C. elegans* vulval induction provides an ideal study system to understand how organismal nutrient status can alter activities of major signaling pathways and penetrance of mutant phenotypes.

A potentially important factor that could partly explain observed starvation effects on vulval induction is prolonged developmental time cause by either starvation exposure and mutation of *pept-1* and *rsks-1*. Applied starvation treatments caused a strong delay in development: animals remained in the L2 stage throughout the 48-hr starvation period (the generation time under normal conditions at 20° is 3.5 days); and *pept-1* and *rsks-1* mutants also showed important developmental delays relative to the wild type. In addition, we observed that individuals starved for only short time period (2 hours) showed no suppression while longer starvation periods (19-24 hours versus 48-120 hours) had different effects, with prolonged starvation periods causing increased suppression (Figure 2B). If the vulval signaling network does not operate at steady state, such developmental delays may allow for accumulation of active downstream effectors inducing vulval fates. Previous experiments, using hydroxyurea treatment or anchor cell ablation, supported this hypothesis (Canevascini et al. 2005; Félix 2007; Wang and Sternberg 1999). However, contrary to this hypothesis, slowing developmental speed by lowering growth temperature does not increase vulval induction of *lin-3(rf)* mutants (Braendle and Félix 2008). Moreover, a mutation in *clk-1* (a gene necessary for the synthesis of ubiquinone) that causes a 2-fold increase in developmental time (Ewbank et al. 1997; Shibata et al. 2003) does not increase vulval induction in a *lin-3(n378)* background (data not shown). Thus, a prolonged developmental time alone may not be sufficient to affect vulval induction.

Our results clarify some seemingly contradictory studies reporting both positive (Braendle and Félix 2008; Ferguson and Horvitz 1985) and negative effects (Battu et al. 2003) of starvation on vulval induction. We confirm here that starvation results in a strong and consistent increase in vulval induction, e.g. illustrated by the strong starvation suppression of *lin-3(rf)* mutations (Figure 2). This positive starvation effect appears to act independently of previously reported transmission of negative starvation effects via specific chemosensory components, Insulin or TGF-*β* signaling (Battu et al. 2003). The study by Battu et al. (2003) used sensitized backgrounds causing vulval hyperinduction (Multivulva phenotype), such as a gain-of-function mutation of *let-60/Ras*, to assess starvation effects: starvation decreased vulval induction in strains with overactivated LET-60 and MPK-1/MAPK signaling, yet had no effect on strains producing excessive LIN-3 or LET-23 (Battu et al. 2003). These results were interpreted to suggest that overstimulation of EGF/EGFR may overcome the negative starvation signal, however, it remained ambiguous at which level of the EGF-Ras-MAPK cascade the signal may integrate. The negative starvation effects on strains with excessive LET-60 or MPK-1 activation were abolished in chemosensory-defective mutants (*osm-5, che-3, sra-13*), implying starvation signal transduction via the sensory system (Battu et al. 2003). Moreover, *daf-2(rf)* suppression of the Multivulva phenotype caused by *let-60/Ras(gf)* in food conditions was interpreted to suggest mimicking (of observed) starvation effects via reduced Insulin signaling (Battu et al. 2003), but it was not tested how vulval induction of *daf-2(rf); let-60(gf)* responds to starvation. Our results reporting an overall positive effect of starvation on vulval induction suggest that sensory perception is not required for this effect to occur. Importantly, however, and in line with the results by Battu et al. (2003), we also find that a compromised DAF-2 activity reduces vulval induction in *lin-3(rf)* to similar extents in both food and starvation conditions, yet without abolishing starvation suppression of *lin-3/egf(rf).* Together, these different results suggest that starvation has both positive and negative effects on *C. elegans* vulval induction, with a positive starvation signal acting independently of external sensory inputs, likely mediated by nutrient sensing via TOR-S6K (this study), and a sensory-system-mediated negative starvation signal, likely involving DAF-2 signaling (this study; Battu et al. 2003). Despite the presence of two antagonistic starvation signals acting in parallel, there is a strong net positive starvation effect, suggesting that negative starvation effects are significantly weaker.

Our study and several others (Battu et al. 2003; Braendle et al. 2010; Ferguson and Horvitz 1985; Moghal et al. 2003) show that *C. elegans* vulval cell fate patterning is sensitive to diverse environmental inputs, which may act through distinct mechanisms to affect vulval signaling pathways. While these results show that specific developmental processes and underlying genetic pathways are responsive to the environment, several key questions remain: What are the consequences of this environmental sensitivity? How does the observed modulation of vulval signaling impact function and precision of this patterning process? Fundamentally, two opposed hypothetical scenarios can be considered to address these questions. First, environmental sensitivity of vulval development may be inevitable, so that environmental effects represent inherent environmental sensitivity of involved mechanisms. If this scenario holds true, the question is whether such environmental sensitivity translates into deleterious effects, such as patterning defects. In a second scenario, environmental sensitivity may reflect a specific developmental modulation to maintain or to enhance functioning and precision in different environmental conditions. This scenario would imply a vulval developmental system whose environmental flexibility is partly of adaptive origin. It is currently not known which of these scenarios apply to the observed environmental sensitivity of *C. elegans* vulval development. However, a previous study (Braendle and Félix 2008) has quantified functioning and precision of vulval cell fate patterning in different environmental conditions: wild type animals of multiple *C. elegans* isolates showed a very low rate of patterning defects in diverse, harsh environments, including starvation. Although starvation significantly increased levels of vulval induction (Braendle and Félix 2008), vulval patterning errors indicative of such increased inductive levels (e.g. hyperinduction) remained very rare (N2 strain, starvation: 3/1000 versus control: 2/1000 individuals) (Braendle and Félix 2008). Therefore, this type of starvation modulation of vulval signaling pathways does not translate into an increased rate of corresponding errors, indicating that this process tolerates considerable changes in pathway activities. These observations reinforce the notion that the *C. elegans* vulval developmental system is robust to extensive signal fluctuations of involved pathways (Barkoulas et al. 2013; Braendle et al. 2010; Félix and Barkoulas 2012; Hoyos et al. 2011; Milloz et al. 2008). Nevertheless, although current results clearly demonstrate robustness, i.e. tolerance of the vulval developmental system to environmental variation, it remains to be evaluated whether specific environmental modulation of signaling pathways enhances the fidelity and precision of the vulval patterning output.

## Acknowledgements

We thank Marie-Anne Félix and Joao Picao-Osorio for discussion and comments on previous versions of the manuscript. Strains were provided by the lab of Marie-Anne Félix and the CGC, which is funded by NIH Office of Research Infrastructure Programs (P40 OD010440). CB acknowledges financial support by the Fondation ARC pour la Recherche sur le Cancer and the Centre National de la Recherche Scientifique (CNRS). SG was supported by fellowships from the Ministère de l’ Enseignement Supérieur et de la Recherche and the Fondation ARC pour la Recherche sur le Cancer. AMVV was supported by the French government through ANR10-LABX-54 MEMOLIFE.

